# Intranasal vaccination induced cross-protective secretory IgA antibodies against SARS-CoV-2 variants with reducing the potential risk of lung eosinophilic immunopathology

**DOI:** 10.1101/2022.05.24.493348

**Authors:** Takuya Hemmi, Akira Ainai, Takao Hashiguchi, Minoru Tobiume, Takayuki Kanno, Naoko Iwata-Yoshikawa, Shun Iida, Yuko Sato, Sho Miyamoto, Akira Ueno, Kaori Sano, Shinji Saito, Nozomi Shiwa-Sudo, Noriyo Nagata, Koji Tamura, Ryosuke Suzuki, Hideki Hasegawa, Tadaki Suzuki

## Abstract

To control the coronavirus disease 2019 (COVID-19) pandemic, there is a need to develop vaccines to prevent infection with severe acute respiratory syndrome coronavirus 2 (SARS-CoV-2) variants. One candidate is a nasal vaccine capable of inducing secretory IgA antibodies in the mucosa of the upper respiratory tract, the initial site of infection. However, regarding the development of COVID-19 vaccines, there is concern about the potential risk of inducing lung eosinophilic immunopathology as a vaccine-associated enhanced respiratory disease as a result of the T helper 2 (Th2)-dominant adaptive immune response. In this study, we investigated the protective effect against virus infection induced by intranasal vaccination of recombinant trimeric spike protein derived from SARS-CoV-2 adjuvanted with CpG oligonucleotides, ODN2006, in mouse model. The intranasal vaccine combined with ODN2006 successfully induced not only systemic spike-specific IgG antibodies, but also secretory IgA antibodies in the nasal mucosa. Secretory IgA antibodies showed high protective ability against SARS-CoV-2 variants (Alpha, Beta and Gamma variants) compared to IgG antibodies in the serum. The nasal vaccine of this formulation induced a high number of IFN-γ-secreting cells in the draining cervical lymph nodes and a lower spike-specific IgG1/IgG2a ratio compared to that of subcutaneous vaccination with alum as a typical Th2 adjuvant. These features are consistent with the induction of the Th1 adaptive immune response. In addition, mice intranasally vaccinated with ODN2006 showed less lung eosinophilic immunopathology after viral challenge than mice subcutaneously vaccinated with alum adjuvant. Our findings indicate that intranasal vaccine adjuvanted with ODN2006 could be a candidate that can prevent the infection of antigenically different variant viruses, reducing the risk of vaccine-associated enhanced respiratory disease.

## 1. Introduction

The coronavirus disease 2019 (COVID-19) caused by severe acute respiratory syndrome coronavirus 2 (SARS-CoV-2) has spread worldwide since December 2019 and is presently a major public health concern [1, 2]. Vaccines could be the most promising approach to protect us from the threat of this infectious disease. In fact, several new vaccines have already been in use, showing high vaccine effectiveness [3–6]. Typical examples include the mRNA vaccines [3, 4] and the viral vector vaccines [5, 6]. Particularly in the UK, it is noteworthy that mRNA vaccine, BNT162b2 (Pfizer-BioNTech), was approved for practical use on December 2, 2020, less than a year after the epidemic began [7]. The advantage of the mRNA vaccine is that if the formulation of the vaccine has already been determined, a new vaccine can be launched for practical use in the shortest time, as long as the nucleic acid information of the target antigen is available. However, the long-term adverse events associated with the new modality of vaccines approved for emergency use are currently unknown; therefore, this issue should be adequately evaluated in the future. On the other hand, although the development of inactivated virus vaccines and subunit vaccines as conventional vaccine formulations has been promoted, the development tends to be delayed because of time taken to prepare the antigen. Unlike vaccines of new modality, these conventional vaccines may have the advantage of being less burdensome to the vaccinees because reactogenicity can be assumed to some extent.

Currently, most vaccines are administered intramuscularly or subcutaneously, resulting in the induction of systemic antigen-specific IgG antibodies. In studies on nasal influenza vaccines, we have already shown that a systemic antibody response is effective for reducing mortality and morbidity associated with influenza but insufficient to prevent infection, and that mucosal secretory IgA antibodies induced by intranasal vaccination are highly protective against not only the vaccine-homologous virus but also antigenically different viruses from the vaccine antigen [8–13]. The World Health Organization (WHO) has recommended the development of a COVID-19 vaccine to prevent infection with SARS-CoV-2 variants [14], therefore the study of vaccines inducing mucosal immunity should be accelerated. In fact, 11 candidate intranasal vaccines against SARS-CoV-2 have already been in clinical trials as of May 13, 2022 [15, 16].

However, there may be a risk of lung eosinophilic immunopathology caused by viral infection among COVID-19 vaccinators, a phenomenon known as vaccine-associated enhanced respiratory disease (VAERD) [17–23]. This phenomenon was first observed in the 1960s in a clinical trial of formalin-inactivated respiratory syncytial virus (FI-RSV) and measles vaccines. Two children died of severe pneumonia with eosinophilic infiltration due to natural infection after vaccination in the clinical trial of the FI-RSV vaccine [24]; thus, VAERD cannot be ignored in vaccine studies. Histological analysis of postmortem lung sections revealed immune complex formation and complement activation in the smaller airways. Similar eosinophilic immunopathology was observed in a mouse experiment with SARS-CoV and Middle East respiratory syndrome (MERS)-CoV vaccines. Some studies have suggested that lung eosinophilic immunopathology is due to the induction of T helper 2 (Th2)-shifted immune responses with high levels of antibody responses and insufficient neutralizing ability [21–23]. Although several studies on nasal COVID-19 vaccines in mouse models have already been reported, these studies did not analyze vaccine-induced eosinophilic immunopathology [25, 26]. Using a mouse model, we recently revealed that the immunopathology of pneumonia with eosinophilic infiltration was induced by the infection with SARS-CoV-2 among mice immunized with recombinant spike (S) protein of the virus combined with alum adjuvant [23]. Thus, the U.S. Food and Drug Administration (FDA) recommended that the balance of T-cell responses should be properly assessed to avoid the risk of VAERD for vaccine candidates to be practically used in the future[27].

In this study, we used a mouse model to evaluate both the protective effect and neutralizing antibodies induced by intranasal administration of recombinant trimeric spike protein derived from SARS-CoV-2 combined with synthetic oligodeoxynucleotides containing the CpG motif, ODN2006 [28]. In addition, T-cell responses and the risk of VAERD were examined after viral challenge in immunized mice. Our data showed that intranasal administration of recombinant spike protein with ODN2006, rather than subcutaneous administration in the presence of alum adjuvant, induced neutralizing antibodies with well-balanced T-cell responses, resulting in the protection against homologous or heterologous virus infection without lung eosinophilic immunopathology.

## 2. Materials and Methods

### 2.1. Purification of recombinant SARS-CoV-2 Spike protein

Recombinant trimeric ectodomain of SARS-CoV-2 S protein with two proline mutations (rtS-ecto2P) as a vaccine antigen was produced using a *Drosophila* expression system (Thermo Fisher Scientific, Grand Island, NY). The protein sequence was modified to remove the furin cleavage site (RRAR to GSAG), and two stabilizing mutations were introduced (K986P and V987P; wild-type numbering) [29, 30]. Protein expression and purification were performed as previously described [31]. Recombinant trimeric ectodomain of S protein with six proline mutations (rtS-ecto6P) for enzyme-linked immunosorbent assay (ELISA) was produced using Expi293F cells (Thermo Fisher Scientific). In addition to the mutation at the furin cleavage site, six stabilizing mutations were introduced (F817P, A892P, A899P, A942P, K986P, and V987P; wild-type numbering) [32].

### 2.2. Immunization and sampling

Female BALB/c mice (20-24 weeks old) (Japan SLC Inc., Hamamatsu, Shizuoka, Japan) were maintained in specific pathogen-free facilities. The mice were either intranasally or subcutaneously vaccinated with 3 µg rtS-ecto2P three times at 2-week intervals. Intranasal vaccination was performed by instillation of 6 µL of vaccine solution with or without 10 µg of ODN2006 into each nostril (total, 12 µL/mouse). Subcutaneous vaccination was performed by inoculation with 100 µL of vaccine solution containing 10 µg of ODN2006 (InvivoGen, San Diego, CA) or 1 mg of Imject Alum adjuvant (Thermo Fisher Scientific) in the dorsal part of the cervical region. ODN2006 and Imject Alum were used as adjuvants to induce Th1- and Th2-dominant immune responses, respectively. For the time-course evaluation of the antibody response, partial blood sampling from the orbit was performed at 2-week intervals from the final vaccination. Serum, nasal and lung wash, and cervical lymph nodes were collected from mice for the evaluation of antibody and cellular immune responses, respectively, one week after the final vaccination.

All immunizations and partial blood sampling were performed under anesthesia. All animal experiments were performed in accordance with the Guide for Animal Experiments Performed at the National Institute of Infectious Diseases (NIID) and were approved by the Animal Care and Use Committee of NIID.

### 2.3 Virus challenge and sampling

To evaluate effectiveness of protection against virus infection, vaccinated mice were inoculated intranasally with the mouse adapted SARS-CoV-2 strain, QHmusX (GenBank Accession No.: LC605054) [23], into the lungs (40 LD_50_ per mouse) and nasal cavity (6 LD_50_ per mouse) 2 weeks after the final immunization. To protect against SARS-CoV-2 variants—Alpha variant QHN001 (lineage B.1.1.7, GISAID: EPI_ISL_804007), Beta variant TY8-612 (lineage B.1.351, GISAID: EPI_ISL_1123289), and Gamma variant TY7-501 (lineage P.1, GISAID: EPI_ISL_833366)—were intranasally challenged with 3.5×10^5^ TCID_50_ into the lungs and nasal cavity. Intranasal challenge into the upper and lower respiratory tracts was performed by instillation of 30 µL and 4 µL (2 µL in each nostril), respectively. To determine the viral titer in the nasal mucosa and lungs, nasal and lung washes were collected 3 days after virus challenge, and the lungs were collected for the evaluation of eosinophilic immunopathology at 6 days post-infection. In addition, body weight was monitored for 10 days after the virus challenge. The humane endpoint was defined as the appearance of clinical diagnostic signs of respiratory stress, including respiratory distress and > 25% weight loss. The SARS-CoV-2 challenge was performed in a biosafety level 3 facility according to the Guidelines for Animal Experiments performed at NIID.

### 2.4. Estimation of SARS-CoV-2 S-specific antibody responses

SARS-CoV-2 S-specific antibodies were estimated using ELISA. Half-area flat-bottomed microtiter plates (Corning Inc., NY) were coated with 50 ng/well rtS-ecto6P, followed by blocking with PBS containing 5% skim milk and 0.05% Tween 20. Serial dilutions of serum samples from vaccinated mice were added to each well of microtiter plates. IgG antibodies were detected using biotin-conjugated goat anti-mouse IgG antibody (Jackson Immunoresearch, West Grove, PA), followed by alkaline phosphatase-conjugated streptavidin (Invitrogen, CA, USA). The enzymatic reaction was initiated by the addition of the substrate *p*-nitrophenylphosphate (Sigma-Aldrich, Burlington, MA). The absorbance at 405 nm was measured using an iMark microplate reader (Bio-Rad, Hercules, CA). All procedures were performed at room temperature. The S-specific IgG antibody titer was defined as the reciprocal of the highest dilution of the test sample, giving a higher absorbance than the cut-off value obtained as 2-fold mean absorbance of serial dilutions of control naive mouse serum set in each plate.

Quantification of S-specific IgG1 or IgG2a antibodies in the serum and IgA antibodies in nasal or lung washes was performed as previously described [23]. Chimeric human-mouse monoclonal IgG1, IgG2a, and IgA antibodies bearing variable regions of the S-specific human monoclonal antibody S309 [33] were used as standard antibodies for quantification. Horseradish peroxidase (HRP)-conjugated polyclonal anti-mouse IgG1 antibody (Bethyl Laboratories, Montgomery, TX), anti-mouse IgG2a antibody (Bethyl Laboratories), or polyclonal anti-mouse IgA antibody (Bethyl Laboratories) were used as detection antibodies. The enzymatic reaction was obtained by adding ABTS substrate (Roche, Basel, Switzerland), and the absorbance of 405 nm was measured.

### 2.5. SARS-CoV-2 neutralization assay

The neutralization assay was performed as previously described [23, 34]. Briefly, 50uL of QHmusX (100TCID50) and 50uL of heat-inactivated serum serially diluted by two-fold were mixed and incubated in 96-well microtiter plates for 1h at 37□, followed by the addition of 100 µL of VeroE6-TMPRSS2 cells (JCRB1819, Japanese Collection of Research Bioresources Cell Bank) [35, 36]. After five days of cultivation, samples were examined for viral cytopathic effects (CPEs). Neutralizing antibody titers were determined as the reciprocal of the highest dilution rate at which no CPEs were observed. The neutralization assay was performed in a biosafety level 3 laboratory at the NIID, Japan.

### 2.6. Enzyme-linked immunospot assay

Cells secreting interferon (IFN)-γ, interleukin (IL)-4, or IL-5 were determined using a mouse enzyme-linked immunospot (ELISpot) assay kit (Mabtech, Cincinnati, OH) according to the manufacturer’s instructions. Briefly, in plates pre-coated with anti-mouse IFN-γ, IL-4, or IL-5 antibodies, 3 × 10^5^ cells harvested from the spleen or cervical lymph nodes were incubated for 16 h in the presence of a peptide pool derived from the S protein of SARS-CoV-2 (a mixture of PepTivator SARS-CoV-2 Prot_S, S1, and S+; Miltenyi Biotec, Bergisch Gladbach, Germany). After washing the cells with PBS, biotin-conjugated anti-IFN-γ, IL-4, or IL-5 detection antibodies were added and incubated at RT for 2 h, followed by incubation with ALP-conjugated streptavidin at RT for 1 h. The enzymatic reaction was initiated by addition of BCIP/NBT. Each experiment was performed in duplicate. Spots formed by cytokine-secreting cells were counted and analyzed using ELISpot reader S6 Universal with ImmunoSpot 7.0 software (Cellular Technology, Ltd., Shaker Heights, OH).

### 2.7. Flow cytometric analysis

T follicular helper (Tfh) cells, germinal center B (GCB) cells, and eosinophils were evaluated by flow cytometry. Single cell suspensions were obtained from the cervical lymph nodes, spleen and lungs of immunized mice. One million cells were stained with FVD506 (Thermo Fisher Scientific) for dead cell removal and blocked with anti-mouse CD16/CD32 monoclonal antibody (BD Pharmingen, San Jose, CA). Cell surface markers of Tfh cells were defined as CD4^+^ CD8^-^ PD-1^+^ CXCR5^+^ among TER119^-^ Ly-6G/Ly-6C^-^ CD11b^-^ CD19^-^ populations, and those of GCB cells were defined as CD19^+^ GL7^+^ CD95^+^cells among TER119^-^ Ly-6G/Ly-6C^-^ CD11b^-^ CD3^-^ populations [37]. Eosinophils are defined as CD45^+^ CD11b^+^ CD11c^-^ Ly6G^+^ Siglec-F^-^ cells [38]. The antibodies used in flow cytometric analysis are summarized in Supplementary Table 1. Samples were analyzed with CantoII (BD Biosciences), and data were analyzed using FlowJo software version 10.8.0 (Tree Star Inc., Ashland, OR).

### 2.8. Quantification of SARS-CoV-2 subgenomic RNA

Total RNA was extracted from 125 µL of nasal or lung wash using ISOGEN-LS (Nippon gene, Toyko, Japan) and purified using a Maxwell RSC 48 Instrument (Promega, Madison, WI) with a Maxwell RSC miRNA Plasma and Serum Kit (Promega). Quantification of subgenomic RNA was performed by real-time reverse transcription PCR (RT-PCR) using a QuantiTect Probe RT-PCR Kit (QIAGEN, Hilden, Germany) with primers and probes as previously described [39]. Real-time RT-PCR was performed using Mx3005P (Stratagene, La Jolla, CA, USA).

### 2.9. Immunohistochemistry

The lungs collected from mice were fixed in phosphate buffer containing 10% formalin. The fixed lungs were embedded in paraffin and sectioned. Eosinophils were identified using Astra Blue/Vital New Red staining (C.E.M. Stain Kit, Diagnostic Biosystems, Pleasanton, CA). The lung tissue sections were observed for eosinophil infiltration in the peribronchiolar areas using an optical microscope.

### 2.10. Statistical analysis

Data analysis and visualization were performed using GraphPad Prism 7.0 software (GraphPad Software Inc., San Diego, CA). For statistical analysis, the Kruskal-Wallis test with Dunn’s multiple comparison test was used for comparisons between groups. Comparison of body weight and survival was performed using Dunnett’s multiple comparisons test following the mixed-effects model or log-rank (Mantel-Cox) test, respectively. Statistical significance was set at P < 0.05.

## 3. Results

### 3.1. Intranasal vaccines induced S-specific antibodies in nasal mucosa and lung as well as serum

To evaluate the S-specific antibody responses, serum, nasal and lung wash specimens were collected from mice that received three doses of either intranasal or subcutaneous vaccines (Fig. 1A). Intranasal vaccination was performed with or without ODN2006 as a mucosal adjuvant and subcutaneous vaccination was performed with ODN2006 or alum adjuvant. Naïve mice were used as negative controls. Serum S-specific IgG antibody titers and neutralization titers were determined using ELISA and microneutralization assays, respectively. The concentration of S-specific IgA antibodies in the nasal or lung wash samples was quantified using ELISA. Intranasal administration without mucosal adjuvant induced low levels of serum S-specific IgG antibodies. In contrast, intranasal vaccination adjuvanted with ODN2006 successfully induced S-specific IgG antibodies in serum at a level similar to that induced by subcutaneous vaccination in the presence of ODN2006 or alum adjuvant (Fig. 1B). The results for the neutralizing antibody titer were similar to those of the S-specific IgG antibody titer (Fig. 1C). Although intranasal vaccination in the absence of mucosal adjuvant failed to induce local antibody responses, nasal and lung S-specific IgA antibodies were detected among intranasally immunized mice in the presence of ODN2006 (Fig. 1D and 1E). No secretory IgA antibodies were detected in samples from subcutaneously vaccinated mice.

**Fig. 1.**
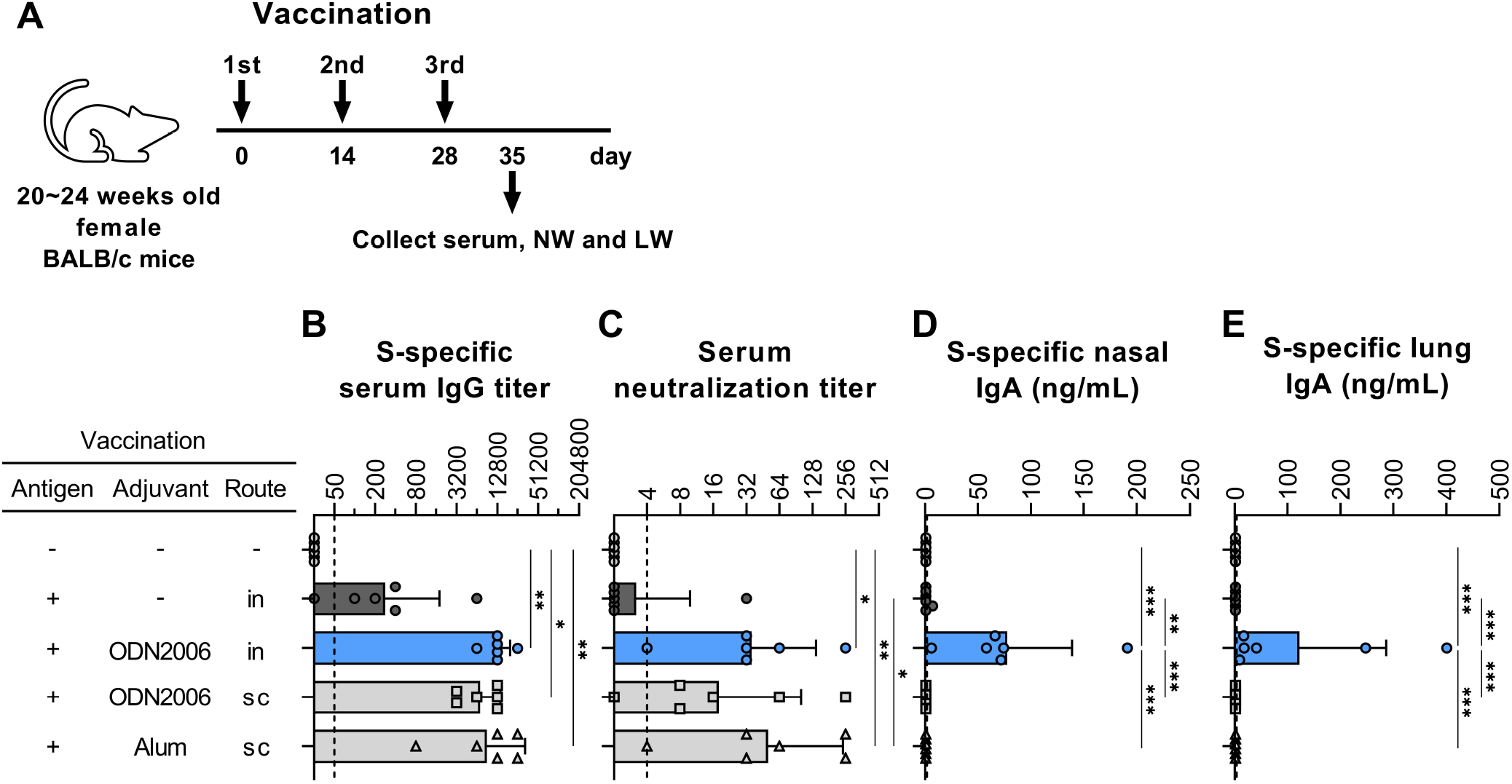
The induction of S-specific antibodies in serum, nasal mucosa and lungs of mice vaccinated intranasally. (**A**) Each of six mice were vaccinated three times at 2-week intervals. One week after the final vaccination, serum, nasal wash (NW) and lung wash (LW) were collected for the evaluation of antibody responses. (**B**, **C**) SARS-CoV-2 S-specific IgG titers and neutralizing antibody titers in sera were measured by ELISA and microneutralization assay, respectively. Graphs shown as the geometric mean titers ± the geometric standard deviation (SD). The dashed line indicates the detection limit of measurement. (**D**, **E**) The concentration of S-specific IgA antibodies in NW and LW was estimated by ELISA. Data shown as the means ± SD. Each dotted line indicates the detection limit of measurement. The p-values were calculated by Kruskal-Wallis test followed by Dunn’s multiple comparison test (*P < 0.05, **P < 0.01). in: intranasally, sc: subcutaneously.

These results indicated that intranasal vaccination with ODN2006 as a mucosal adjuvant induced not only secretory IgA antibodies in the nasal mucosa and lungs but also systemic IgG antibodies at the same level as those obtained from subcutaneous vaccination with ODN2006 or alum adjuvant.

### 3.2. Intranasal vaccine adjuvanted with ODN2006 effectively protect mice from SARS-CoV-2 infection

To evaluate the protection against viral challenge, intranasal inoculation of the mouse-adapted SARS-CoV-2 strain, QHmusX, was performed on mice vaccinated under the same conditions as described above (Fig. 2A). Nasal and lung wash samples were collected at three days post-infection (dpi), and the amount of subgenomic RNA (sgRNA) derived from SARS-CoV-2 in these samples was evaluated by real-time RT-PCR to assess the protection of mice against infection. The number of sgRNA copies in the nasal wash was significantly reduced in mice with S-specific IgA in the nasal mucosa induced by intranasal vaccination adjuvanted with ODN2006 (Fig. 1D and 2B). No significant decrease in sgRNA in the nasal mucosa was observed in mice that received subcutaneous vaccine or mice intranasally vaccinated with antigen only. A significant decrease in sgRNA copies in the lung wash was observed in mice vaccinated intranasally or subcutaneously in the presence of ODN2006 or alum, respectively, which showed high neutralizing antibody titers in serum (Fig. 1C and 2C). Simultaneously, the mice were monitored for body weight and survival for 10 days after the challenge (Fig. 2D and 2E). Naïve mice and those intranasally vaccinated with antigen only died by 6 dpi. Among mice that received subcutaneous vaccine with ODN2006 or alum, although few mice died, others recovered from the apparent decrease in body weight post challenge and survived until 10 dpi; in contrast, all mice intranasally vaccinated with ODN2006 survived without remarkable body weight change during the observation period.

**Fig. 2.**
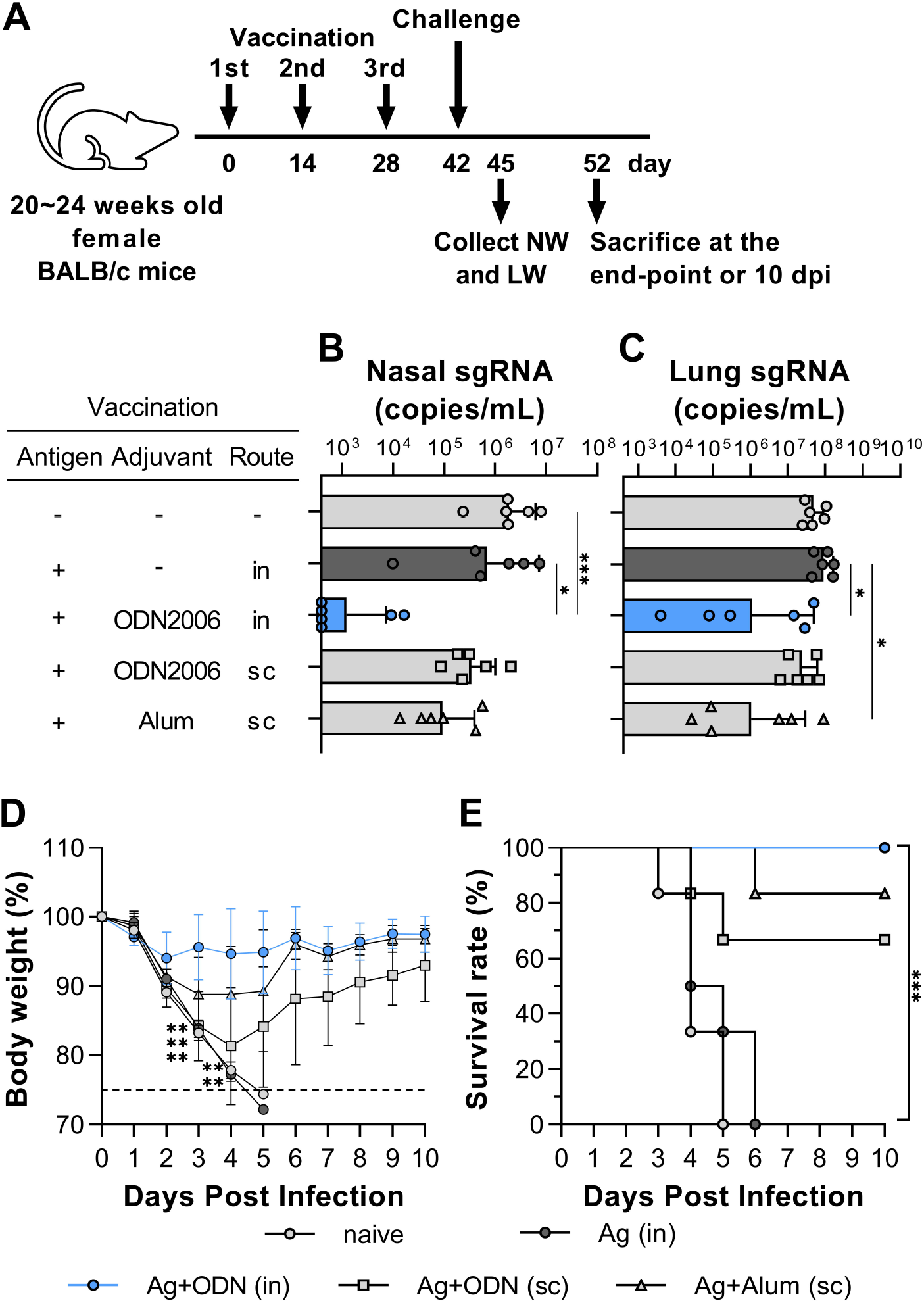
Intranasal vaccination adjuvanted with ODN2006 protected mice from the lethal challenge with SARS-CoV-2. (**A**) Each of 12 mice were vaccinated three times at 2-week intervals. Two weeks after the final vaccination, mice were intranasally challenged with mouse adopted SARS-CoV-2 strain, QHmusX, into both lungs and nasal cavity (40 and 6 LD_50_ per mouse, respectively). NW and LW were collected from each of six mice for the evaluation of antibody responses at 3 dpi. Body weight and survival of six mice were monitored 10 days after viral challenge. (**B**, **C**) Copy numbers of SARS-CoV-2 subgenomic RNA (sgRNA) in NW and LW were evaluated by real-time RT-PCR. Data shown as the geometric mean ± the geometric SD. The p-values were calculated by Kruskal-Wallis test followed by Dunn’s multiple comparison test (*P < 0.05, *** P < 0.001). (**D**, **E**) Body weight changes and survival rates during 10 days of observation after challenge with QHmusX. Data shown as the means ± SD. The p-values of body weight and survival were compared with mice intranasally vaccinated with ODN2006 by mixed-model analysis followed by Dunnett’s multiple comparisons test and log-rank (Mantel-Cox) test (**P < 0.01).

In addition, protection against viral challenge with SARS-CoV-2 variants (Alpha, Beta and Gamma variants) was assessed (Fig. 3A). Significant reductions in nasal sgRNA derived from each variant virus were achieved in mice intranasally vaccinated in the presence of ODN2006 when compared to naïve mice (Fig. 3B–3D). In the lungs, sgRNA of Alpha and Gamma variants significantly decreased by both intranasal and subcutaneous vaccination with antigen together with ODN2006, while that of Beta variant presented a significant decrease only by intranasal but not subcutaneous vaccination (Fig. 3E–3G).

**Fig. 3.**
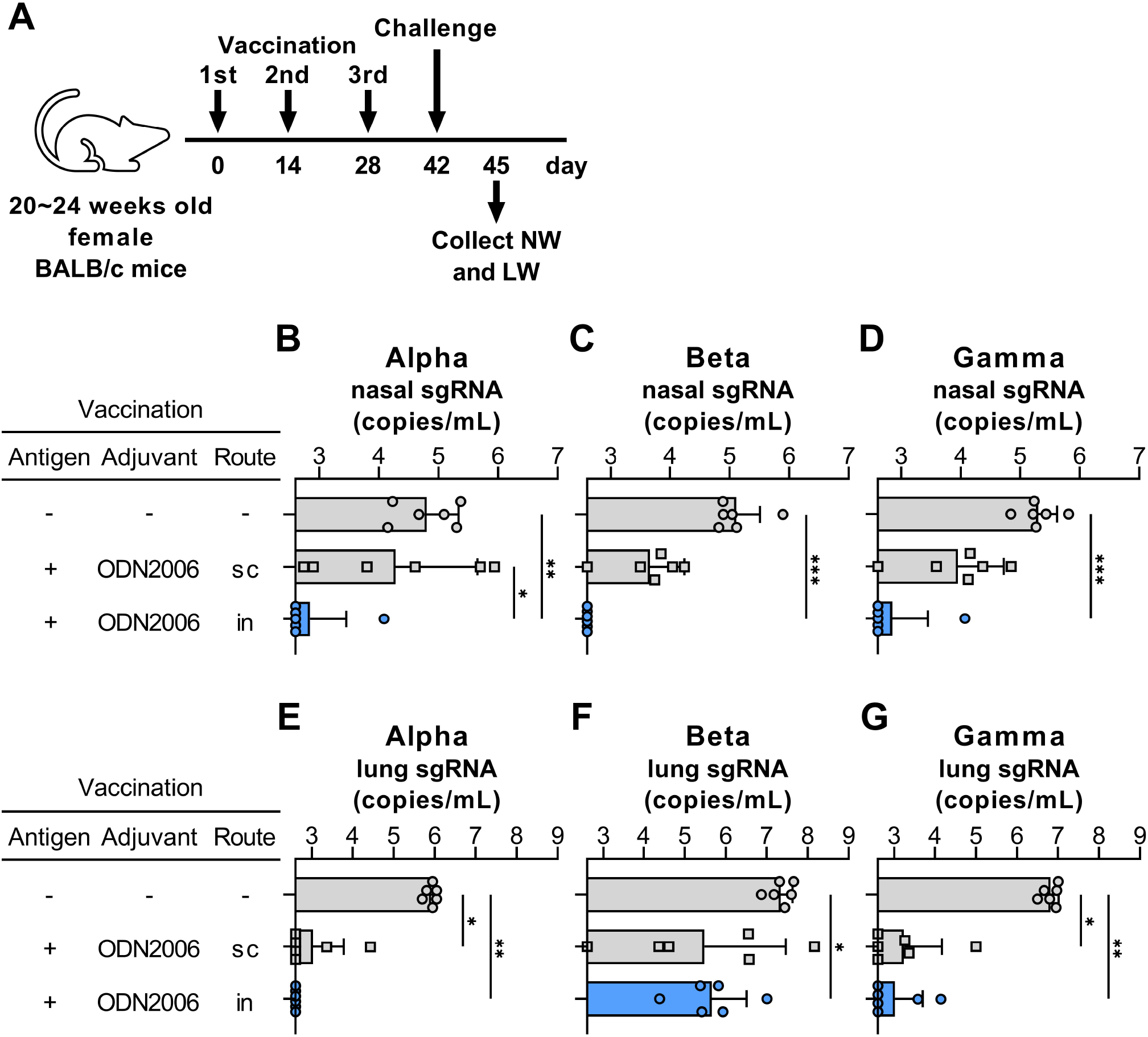
Secretory IgA antibodies were protective against SARS-CoV-2 variants. (**A**) Mice were vaccinated three times at 2-week intervals. Two weeks after the final vaccination, each of six mice were intranasally challenged with 3.5×10^5^ TCID_50_ of Alpha (**B**, **E**), Beta (**C**, **F**) or Gamma (**D**, **G**) variant into both lungs and nasal cavity. At 3 dpi, NW and LW were collected. Copy numbers of sgRNA in NW (**B**-**D**) and LW (**E**-**G**) were evaluated by real-time RT-PCR. Data shown as the geometric mean ± the geometric SD. The p-values were calculated by Kruskal-Wallis test followed by Dunn’s multiple comparison test (*P < 0.05, *** P < 0.001).

These findings indicate that IgA antibodies in the nasal mucosa and lung induced by intranasal, but not subcutaneous, vaccination combined with ODN2006 were cross-protective against SARS-CoV-2 variants.

### 3.3. Formation of germinal center and maintenance of antibody responses by intranasal vaccination

The induction of memory B cells and long-lasting humoral immune responses derived from long-lived plasma cells is important for successful vaccine-evoking protective antibody responses [40]. The formation of a germinal center is required for the affinity maturation of antibodies and determination of the B cell life span [37]. Hence, vaccination-induced changes in the percentage of Tfh cells and GCB cells were evaluated in lymphocytes from draining cervical lymph nodes using flow cytometry. The proportions of both Tfh and GCB cells in cervical lymph nodes were definitely increased by intranasal vaccination with antigen plus ODN2006 (Fig. 4A and 4B). In addition, systemic S-specific IgG antibody responses were evaluated in serum samples collected for 20 weeks at 2-week intervals after the initial vaccination (Fig. 4C). The highest responses were achieved two weeks after the final vaccination, and mice that received subcutaneous vaccine adjuvanted with alum or ODN2006 showed the highest S-specific IgG antibodies, followed by mice intranasally vaccinated in the presence of ODN2006. Although systemic S-specific IgG antibodies slightly declined over time during the observation period, IgG antibodies were well held in mice subcutaneously vaccinated with alum compared to mice that received intranasal or subcutaneous vaccination with ODN2006 (Fig. 4D). At 20 weeks after the initial vaccination, S-specific nasal IgA antibodies were detected in five out of six mice with intranasal vaccination combined with ODN2006 but not in mice with other vaccinations (Fig. 4E).

**Fig. 4.**
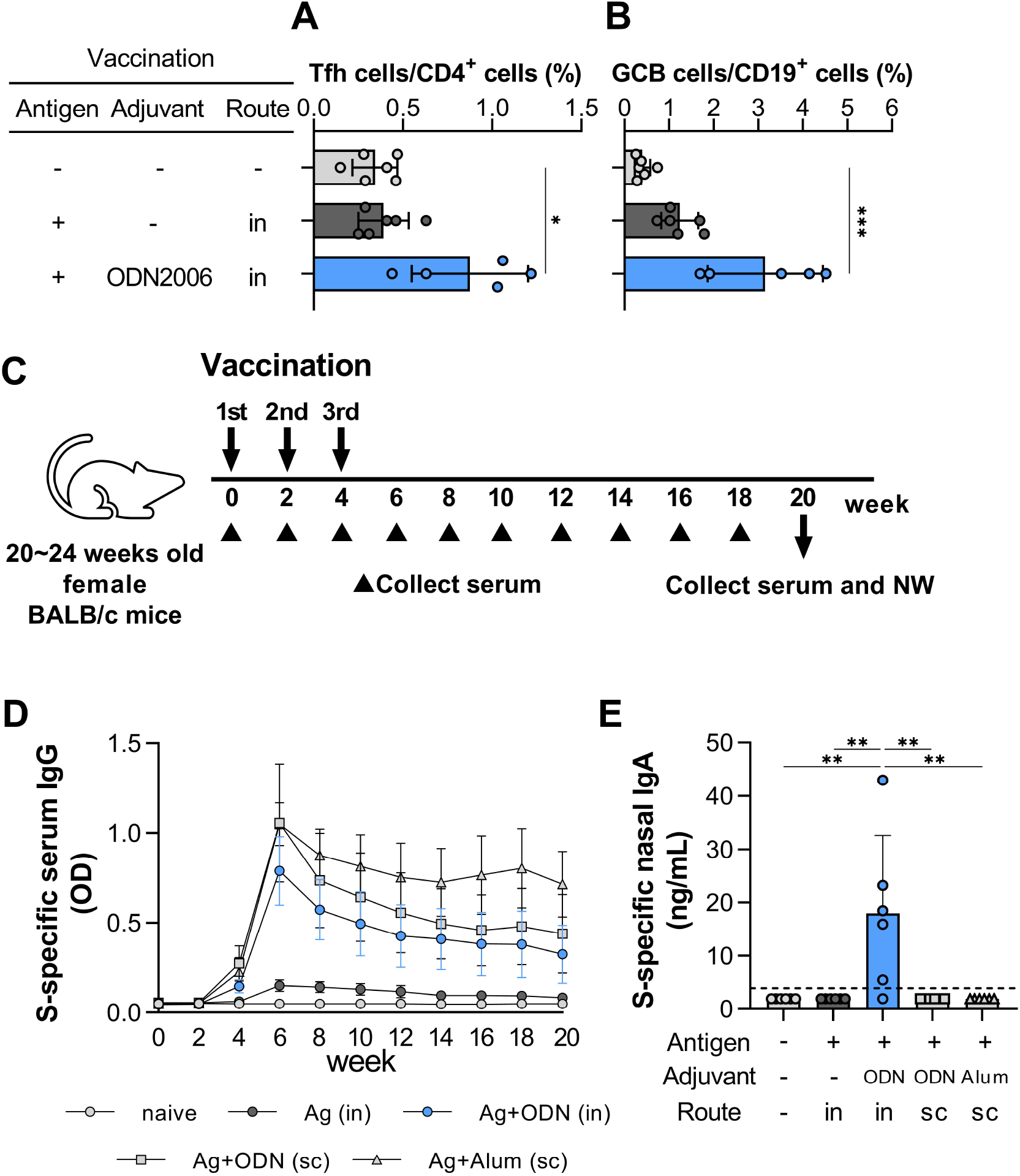
Germinal center formation and long-lasting antibody responses. (**A**, **B**) The frequency of Tfh cells (CXCR5^+^ PD-1^+^) among CD4^+^ cells and GCB cells (GL7^+^ CD95^+^) among CD19^+^ B cells was evaluated by flow cytometry in cervical lymph nodes collected from mice one week after third vaccination. Results were shown as the mean ± SD. (**C**) To evaluate time-course of antibody responses induced by three doses of vaccination, serum samples were collected every two weeks. After 20 weeks from the initial vaccination, serum and NW samples were collected (six mice per group). (**D**) The changes of optical density (405 nm) of S-specific IgG antibodies were shown as the mean ± SEM. (**E**) S-specific IgA concentration in NW collected at 20 weeks after the initial vaccination was evaluated by ELISA. Results were shown as the mean ± SD. Each dotted line indicates the detection limit of measurement. The p-values were calculated by Kruskal-Wallis test followed by Dunn’s multiple comparison test (*P < 0.05, **P < 0.01, ***P < 0.001).

Overall, these results suggest that intranasal vaccination with ODN2006 induced the formation of GCs in the draining lymph nodes and the maintenance of local secretory IgA antibody responses in the nasal mucosa, despite a slight decline in systemic IgG responses.

### 3.4. Induction of Th1 response by using ODN2006 as an adjuvant

Lung eosinophilic immunopathology as a phenomenon of VAERD was observed in a clinical trial of FI-RSV and measles vaccine, and in a mouse model of SARS- or MERS-CoV vaccine. This phenomenon is suspected to be dependent on the Th2 dominant immune response of vaccine candidates [21–23]. Therefore, the Th response induced by intranasal vaccination with recombinant S protein in the presence of ODN2006 should be carefully examined. In mice, Th1 cells produce IFN-γ resulting in IgG2a induction, in contrast, Th2 cells produce IL-4 and IL-5 inducing IgG1 responses [41].

Cells isolated from the spleen and cervical lymph nodes one week after the final immunization were evaluated for cytokine production under stimulation of the peptide pool of S protein by ELISpot assay (Fig. 5A). Significant induction of IFN-γ-secreting cells was observed in the splenocytes of mice subcutaneously vaccinated with ODN2006 (Fig. 5B). In contrast, a significant induction of IL-4-secreting cells was observed in splenocytes obtained from mice subcutaneously vaccinated with alum adjuvant, and a similar tendency was observed in IL-5-secreting cells (Fig. 5C and 5D). In the case of splenocytes, each cytokine-secreting cell significantly increased in mice that received subcutaneous vaccination with ODN2006 or alum but not in those intranasally vaccinated. Therefore, cytokine-secreting cells were evaluated using cells isolated from the draining cervical lymph nodes of intranasally vaccinated or naive mice. Intranasal vaccination in the presence of ODN2006 significantly increased the number of IFN-γ-secreting cells, while IL-4 and IL-5 secreting cells decreased compared to intranasal vaccines with only antigen (Fig. 5E). In contrast, intranasal vaccination with antigen in the absence of mucosal adjuvant significantly induced IL-4 and IL-5 secreting cells, but not IFN-γ-secreting cells, in draining lymph nodes (Fig 5F and 5G).

**Fig. 5.**
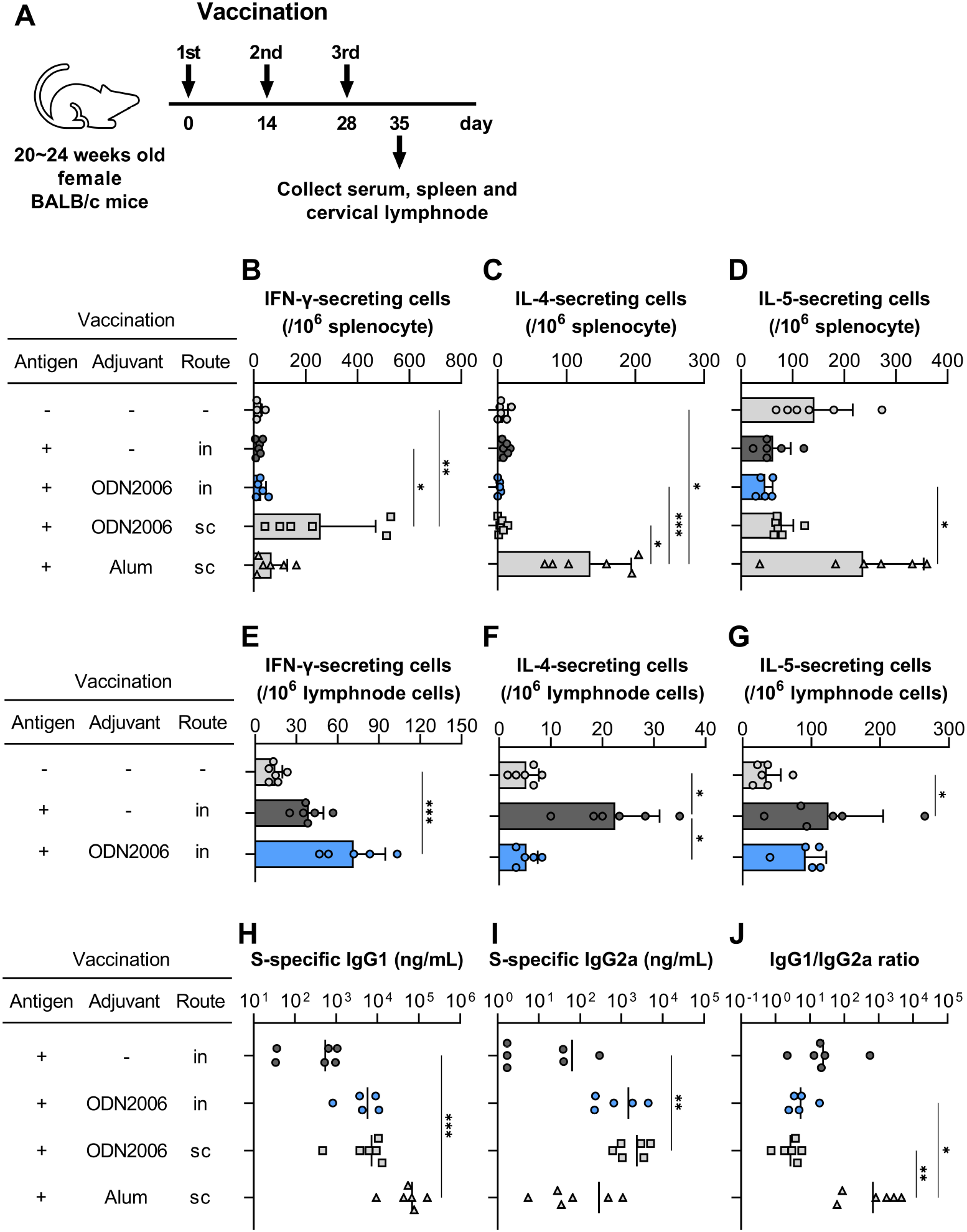
Induction of significant Th1 response by using ODN2006 as an adjuvant. (**A**) Spleen and cervical lymph nodes were collected from individuals shown in Figure.1(**A**). Single cell suspensions obtained from each tissue was cultured under the stimulation of peptide pool of SARS-CoV-2 S protein. Spleen (**B**-**D**) or cervical lymph node cells (**E**-**G**) were counted on the production of IFN-r (**B**, **E**), IL-4 (**C**, **F**) and IL-5 (**D**, **G**) by ELISpot assay. Data shown as the means ± SD. (**H**, **I**) SARS-CoV-2 S-specific IgG1 and IgG2a antibodies were quantified by ELISA, and (**J**) IgG1/IgG2a ratio was calculated. Data shown as the geometric means ± geometric SD. The p-values were calculated by Kruskal-Wallis test followed by Dunn’s multiple comparison test (*P < 0.05, **P < 0.01, ***P < 0.001).

The dominant T cell response was estimated using an S-specific IgG antibody subclass (Fig. 5H and 5I). Large amounts of S-specific IgG1 antibodies were obtained in mice subcutaneously vaccinated with alum adjuvant compared to mice immunized in the presence of ODN2006; however, S-specific IgG2a antibodies in mice intranasally or subcutaneously vaccinated in the presence of ODN2006 were higher than those obtained in mice vaccinated with alum. When the S-specific IgG1/IgG2a ratio was calculated, the IgG1/IgG2a ratio after vaccination with ODN2006 was significantly lower than that after vaccination with alum (Fig. 5J).

These results, obtained from the estimation of cytokine-secreting cells and IgG1/IgG2a ratio, suggested that ODN2006, used as an adjuvant, induced Th1 dominant immune responses regardless of the administration route.

### 3.5. Lung eosinophilic immunopathology is reduced by vaccines that induce a remarkable Th1 response

The correlation between vaccine-induced Th2 dominant immune responses and lung eosinophilic immunopathology was evaluated in a lethal challenge model in mice (Fig. 6A). S-specific IgG1/IgG2a ratios were calculated by ELISA in sera obtained one week prior to challenge, and eosinophilic infiltration into the lungs was examined at 6 dpi by histopathological and flow cytometric analyses. As shown in Fig. 6B, histological analysis revealed that mice immunized in the presence of ODN2006, regardless of the route of vaccination, showed small lesions with infiltration of inflammatory cells, including neutrophils and mononuclear cells around the blood vessels and bronchi, but little eosinophil infiltration. In contrast, eosinophilic infiltration around the bronchi and blood vessels was observed in mice intranasally vaccinated with only antigen or subcutaneously vaccinated with alum. Similar results were obtained in the flow cytometric analysis. Although there were no significant differences in the percentages of eosinophils induced among the different vaccines, these values correlated well with the S-specific IgG1/IgG2a ratios (r = 0.779, p = 0.0015) (Fig. 6C and 6D).

**Fig. 6.**
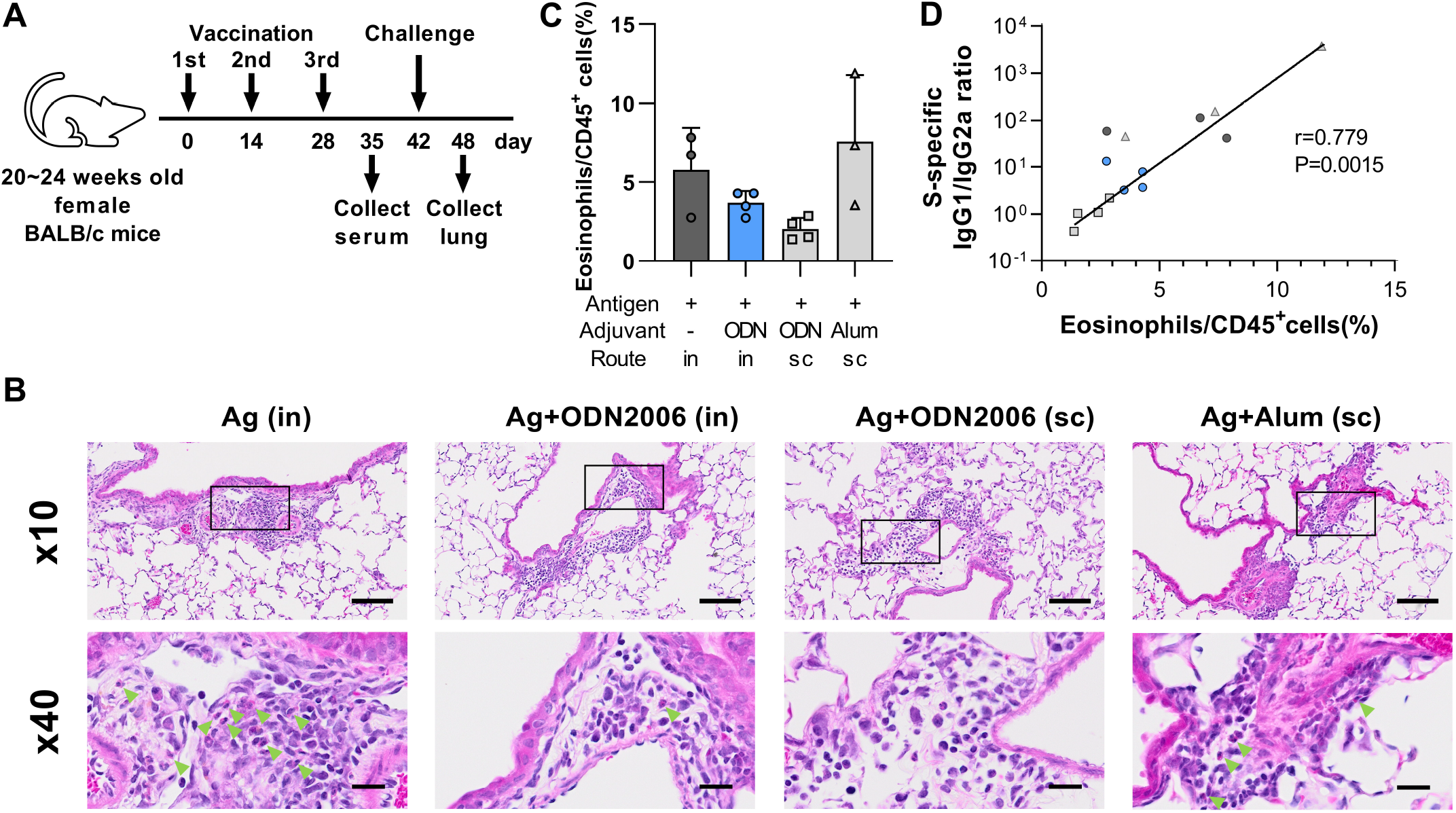
Eosinophilic infiltration into the lungs was suppressed in mice vaccinated with ODN2006 inducing a remarkable Th1 response. (**A**) Each of six mice were vaccinated three times at 2-week intervals. Two weeks after the final vaccination, mice were intranasally challenged with mouse adopted SARS-CoV-2 strain, QHmusX, into both lungs and nasal cavity (40 LD_50_ and 6 LD_50_ per mouse, respectively). At 6 dpi, infiltration of eosinophils was evaluated by histopathological or flow cytometric analysis, on lungs collected from surviving 1∼2 or 2∼4 mice in each group, respectively. (**B**) Histopathological findings of mouse lungs by eosinophil staining using the combined eosinophil-mast cell staining (C.E.M.) kit. Green arrow heads point to eosinophils. The images in the lower panels are enlargements of area boxed in the upper images. The scale bar is 200 µm for low magnification and 20 µm for high magnification. (**C**) The percentage of eosinophils (CD11b^+^ CD11c^-^ Siglec-F^+^ Ly-6G^-^) among CD45+ cells was analyzed by flow cytometry. Results were shown as the mean ± SD. The p-values were calculated by Kruskal-Wallis test followed by Dunn’s multiple comparison test. (**D**) Correlation between S-specific IgG1/IgG2a ratio and the frequency of eosinophils in lung cells was analyzed by Spearman correlation. S-specific IgG1/IgG2a ratio was evaluated using serum samples collected seven days before the virus challenge.

Our results showed that Th2 dominant immune response, suspected by large values of IgG1/IgG2a, caused lung eosinophilic immunopathology. In contrast, in immunization combined with ODN2006, Th1 shifted immune responses correlating with low values of IgG1/IgG2a ratio alleviated the risk of eosinophilic infiltration.

## 4. Discussion

In the current study, we revealed that intranasal vaccination with S protein together with ODN2006, a toll-like receptor 9 agonist [28], could induce cross-protective secretory IgA antibodies against SARS-CoV-2 variants in the nasal mucosa, which is the initial site of infection. Furthermore, we demonstrated that this vaccine could reduce the potential risk of lung eosinophilic immunopathology in the case of post-vaccination infection.

In our previous studies on intranasal influenza vaccine, it has been revealed that intranasal vaccination induces not only IgG antibodies in the serum but also cross-protective secretory IgA antibodies on the surface of mucosal epithelial cells in the upper respiratory tract [8–13]. Here, we evaluated serum and mucosal antibody responses and protective effects induced in mice by intranasal vaccination with recombinant SARS-CoV-2 S protein. Only mice immunized intranasally with antigen combined with ODN2006 induced mucosal IgA as well as systemic IgG antibody responses accompanied by a significant reduction in viral load in both the upper respiratory tract and lungs. All individuals receiving this vaccine survived a lethal challenge with the mouse adapted SARS-CoV-2 strain without significant weight loss. In addition, the cross-protective ability by nasal vaccine was evaluated against SARS-CoV-2 Alpha, Beta or Gamma variant. Results of neutralization assays using human sera collected from mRNA vaccinees or individuals who suffered from breakthrough infections suggest that the antigenicity of the Beta variant differs from those of the Alpha and Gamma variants. [42–44]. When SARS-CoV-2 variants were challenged into the lungs, infections of Alpha or Gamma variants were suppressed in mice intranasally or subcutaneously vaccinated, whereas infection with the Beta variant could not be prevented by either vaccination. On the other hand, all variants challenged into the nasal cavity were significantly prevented in mice possessing mucosal secretory IgA antibody induced by intranasal vaccination, but not in those subcutaneously vaccinated. These results indicate that, compared to systemic IgG antibodies which are primarily responsible for protection against infection in the lungs [11], secretory IgA antibodies that can be induced by intransal vaccination possess higher cross-protective activity against SARS-CoV-2 variantviruses with different antigenicities. Thus, a nasal vaccine that could induce highly cross-protective secretory IgA antibodies in the nasal mucosa, which is the gateway to respiratory infection, would be a more effective, reasonable vaccine candidate. It has been shown that the Omicron variant, which is currently prevalent around the world, could replicate more effectively in the bronchi than in the lungs, compared with other variants and the ancestor [45–48]. Therefore, there is high chance that a nasal vaccine which could induce cross-protective IgA antibodies in the upper respiratory tract would be effective against the Omicron variant as well.

When considering the COVID-19 vaccine, there is a concern about the potential risk of lung eosinophilic immunopathology in post-vaccination infections as VAERD. The FDA recommends addressing the potential risk of VAERD in animal models [27]. Since it has been considered that a Th2-dominant response increases the risk of VAERD, we investigated the T-cell response induced by intranasal vaccination combined with ODN2006 in detail. While IL-4- or IL-5-secreting cells were highly observed in spleen collected from mice subcutaneously vaccinated with alum as a typical Th2 adjuvant, individuals who received subcutaneous vaccination with ODN2006 showed high amounts of IFN-γ-secreting cells. Although nasal vaccination had no significant impact on spleen cells compared to subcutaneous vaccination, IFN-γ-secreting cells were significantly detected in draining cervical lymph nodes of intranasally vaccinated individuals with ODN2006. It was suggested that ELISpot assay using spleen cells might be unsuitable for the evaluation of T cell responses induced by intranasal vaccines. In contrast, the IgG1/IgG2a ratios determined from the quantification of S-specific IgG1 and IgG2 subclasses in serum were likely able to assess T cell responses reflecting neutralizing antibody titers, because the tendency to show similar levels of the IgG1/IgG2a ratio among mice subcutaneously or intranasally vaccinated with ODN2006 was consistent with that of neutralizing antibody titers. Interestingly, eosinophil infiltration into the lung correlated well with the serum IgG1/IgG2a ratio in our mouse model. At this time, although the threshold for eosinophil infiltration that causes eosinophilic pneumonia is unknown, analyzing immunization-induced IgG subclasses could be an indicator for estimating whether there is a potential risk of VAERD. Although several substances have been reported as potential mucosal adjuvants (e.g., cholera toxin B subunit and synthetic double-stranded RNA), the potential risk of lung eosinophilic immunopathology should be adequately investigated when designing a COVID-19 vaccine [8, 13].

In addition, the induction of B-cell memory and long-lived plasma cells by vaccination is noteworthy, since germinal center formation is essential not only for the production of high-affinity antibodies, but also for the determination of the B cell life span.. The induction of Tfh and GCB cells by vaccination is essential to address this issue [47, 49, 50]. In the current study, intranasal vaccination in the presence of ODN2006 successfully induced Tfh and GCB cells, long-lasting IgG antibodies in the serum, and IgA antibodies in the nasal mucosa up to 16 weeks after the final vaccination.

Considering the emergence of new SARS-CoV-2 variants, nasal vaccines inducing a secretory IgA antibody with high cross-protective ability on the mucosal epithelium of the upper respiratory tract, which is the site of infection, could be highly useful, avoiding repeated vaccinations using newly manufactured antigens. In conclusion, our study showed that an intranasal COVID-19 vaccine of recombinant spike protein combined with an adjuvant inducing a Th1-shifted response would be a safe and effective vaccine not only for preparing cross-protective secretory IgA antibodies, but also to reduce the potential risk of VAERD, that is, lung eosinophilic immunopathology.

## Acknowledgements

We thank A. Kojima and A. Sataka (Department of Pathology, NIID) and N. Kurisaki (Department of Virology, Kyushu Univ) for excellent support, and S. Takatsuka (Department of Fungal Infection, NIID) for technical suggestions and helpful discussions. Additionally, we would like to thank Editage (www.editage.com) for English language editing.

## Author Contributions

Conceptualization: A.A, T.He., and T. S

Methodology: T.He., A.A., T.Ha., N.I.-Y., S.I., Y.S. A.U., K.S., N.S.-S., N.N. and T.S.

Investigation: T.He, T.Ha, M.T., T.K., N.I.-Y., Y.S., S.M., A.U., K.S., S.S. and A.A.

Visualization: T.He. and A.A.

Funding acquisition: H.H. and T.S.

Project administration: H.H. and T.S.

Supervision: R.S., K.T., H.H., and T.S.

Writing—Original draft: T.H. and A.A.

Writing—Review and editing: T.He., A.A., T.Ha., M.T., T.K., N.I.-Y., S.I., Y.S., S.M., A.U., K.S., S.S., N.S.-S., N.N., K.T., R.S., H.H. and T.S.

