## Supplementary Table1 for "Intranasal vaccination induced cross-protective secretory IgA antibodies against SARS-CoV-2 variants with reducing the potential risk of lung eosinophilic immunopathology"

**Supplementary Table 1. List of antibodies used for flow cytometry analysis**

| Antibodies | Conjugation | Clone | Source | Catalogue number |
|---|---|---|---|---|
| TER119 | BV510 | TER-119 | BioLegend | 116237 |
| Ly-6G/Ly-6c | BV510 | RB6-8C5 | BioLegend | 108457 |
| CD11b | BV510 | M1/70 | BioLegend | 101263 |
| CD19 | BV510 | 6D5 | BioLegend | 115545 |
| CD4 | Pacific Blue | RM4-5 | BioLegend | 100531 |
| CD8 | AF700 | 53-6.7 | BioLegend | 100730 |
| PD-1 | PE | RMPI-30 | BioLegend | 109103 |
| CXCR5 | APC | L138D7 | BioLegend | 145505 |
| CD3 | BV510 | 145-2C11 | BioLegend | 100353 |
| CD19 | FITC | 1D3 | BioLegend | 152403 |
| GL7 | PE | GL7 | BioLegend | 144607 |
| CD95 | APC | SA367H8 | BioLegend | 152603 |
| CD45 | APC | 30-F11 | BioLegend | 103112 |
| CD11b | AF488 | M1/70 | BioLegend | 101217 |
| CD11c | Pacific Blue | N418 | BioLegend | 117322 |
| Ly-6G | PerCP | 1A8 | BioLegend | 127654 |
| Siglec F | PE | S17007L | BioLegend | 155506 |
